# Rodent chronic variable stress procedures: a disjunction between stress entity and impact on behaviour

**DOI:** 10.1101/2024.07.04.602063

**Authors:** Nicola Romanò, John Menzies

## Abstract

Chronic variable stress (CVS) procedures are widely used to model depression in laboratory rodents. We systematically documented the experimental design used in mouse CVS studies, and the design of the behavioural tests used to evaluate the effect of CVS. In a subset of studies, we measured effect sizes in behavioural tests. Across 202 mouse studies, 82% used a unique CVS procedure. We took advantage of this variability to ask whether the duration and intensity of CVS procedures correlated with effects sizes obtained in five commonly-used behavioural tests: the sucrose preference test (SPT), the tail suspension test (TST), the forced swim test (FST), the open field test (OFT) and the elevated plus maze (EPM). The most evident impact of CVS procedure design on effect sizes were seen in the FST where longer-duration CVS procedures with more diverse types of stressors were associated with a smaller effect size. Next, we correlated effect sizes *between* behavioural tests to explore whether these tests might measure similar or different consequences of CVS. We found a positive correlation between effects sizes in the TST and FST, and in the OFT and EPM, but the two strongest positive correlations were between the EPM and TST, and between the EPM and FST. CVS studies deliberately impose suffering over long periods, and our data raise scientific and ethical questions around the stress procedures used and the behavioural tests used to evaluate them.

## Introduction

Many studies have questioned whether translational studies accurately model human disease or have the power to predict the clinical effectiveness of interventions^1–4^, with inconsistency in experimental design and issues around reproducibility identified as key limitations^2, 4, 5^. Chronic variable stress (CVS) procedures (also called chronic unpredictable stress or chronic mild stress) involve regular and deliberate long-term imposition of different stressors. The aim is to model depression, with outcomes usually evaluated by behavioural testing. Pioneered by Paul Willner, the procedure was based on how a “thoroughly incompetent technician”^6^ might treat animals. For example, using a leaking water bottle that wets the animal’s bedding, or an improper light-dark cycle that disturbs the animal’s circadian biology. The stressors used in these procedures are often described as “micro-stressors”^6^ or “minor-intensity stressors^7^”. However, the severity of the stressors used in current procedures seems to have reverted to those used in Katz’s foundational series of 1981 studies^8–10^. These stressors are well beyond what even the most incompetent technician would impose. They include physical restraint, exposure to cold or heat, removal of food, water or cage bedding, exposure to hypoxia, noise, stroboscopic lighting or predator odours, social isolation or overcrowding, tilting or shaking the home cage, forced swimming, or painful stimuli^11, 12^. Some studies use combined stressors; for example, social isolation at cold temperatures^13^ or shaking the cage in the presence of loud noise^14^. Because suffering is deliberately imposed in these studies, we reasoned that ethical considerations would be of high importance, and researchers would, therefore, use a design consistent with previous studies that enhances the translational value of the study. However, to the best of our knowledge there is little agreement on how to optimise experimental design in CVS studies^15–21^. This means there may be wide variability in the type and amount of stressors used which, in turn, raises questions about their generalisability.

Turning to the behavioural tests that evaluate the effectiveness of a CVS procedure, their reliability and validity have been explored in detail^21–26^, but we wanted to ask whether the outcomes measured in these behavioural tests relate to the properties of the CVS procedure used, and to explore how these behavioural tests relate to each other in terms of what they measure. To do this, we reasoned that if a post-CVS changes in behaviour reflects a biological and/or psychological consequence of CVS, the effect sizes obtained in behavioural tests should depend in some way on the CVS procedure used. Taking the sucrose preference test (SPT) as an example: this test involves giving an animal a choice between two drinking bottles for a defined period, one containing a sucrose solution and one containing water. A reduced preference for sucrose, compared to non-stressed animals, is deemed to reflect anhedonia^27^. A long CVS procedure with a high number and variety of stressors might be expected to result in a higher level of anhedonia than a milder CVS procedure, and this would be detectable as a larger magnitude effect size in the SPT.

Using a similar approach, we also wanted to explore whether different post-CVS behavioural tests measure the same or different consequences of CVS. Take the tail suspension test (TST) and forced swim test (FST) as examples. Both tests involve placing the animal into a situation that it initially wants to escape from (we can infer this from its behaviour^28^), until at some point it becomes immobile. We do not know the animal’s motivations for these behaviours (nor, presumably, does the animal understand the experimenter’s motivations). However, it is generally deemed that an intervention that reduces the time the animal spends immobile reflects an anti-depressive effect^29^, a conclusion based on an assumption that rodent immobility and human depression are biologically equivalent. But is the immobility observed in the TST and the immobility observed in the FST a reflection of the same consequence of CVS? If it is, we could predict that across a series of studies that used both tests to evaluate a given CVS procedures, the effect sizes will correlate between these tests but will not correlate with effect sizes in, for example, the SPT. A very strong positive correlation would not incontrovertibly demonstrate that both immobility tests measure an identical consequence of CVS, but it would demonstrate that these tests do not distinguish between two or more different (putative) immobility-related consequences of CVS. The same question can be asked of the open field test (OFT) and the elevated plus maze (EPM). Both measure whether an animal prefers to be in an area close to a wall or being in an exposed area. Again, we cannot know the animal’s motivations, or whether and how they relate to motivations and behaviours in people with anxiety, but spending less time in the exposed area is deemed to reflect a higher level of anxiety^24^. If these tests both measure the same or very similar aspects of anxiety, the effect sizes should correlate.

Accordingly, we hypothesised that there will be a positive correlation between CVS characteristics and the magnitude of effect sizes in evaluative behavioural tests. In other words, that a long CVS procedure with a large number of diverse stressors will be associated with large effect sizes in behavioural tests, and that a shorter CVS procedure with fewer stressors will be associated with smaller effect sizes. We also hypothesised that effect sizes in behavioural tests will positively correlate in a way that reflects what they measure. Specifically, that TST and FST effect sizes will positively correlate, and that OFT and EPM effect sizes will positively correlate.

To address this, we systematically documented certain characteristics of CVS procedures, and the types and combinations of the behavioural tests used to evaluate the effect of CVS. In a subset of articles, we also measured effect sizes obtained in the SPT, the TST, the FST, the OFT and the EPM. To address the first hypothesis, we correlated effect sizes in these tests with properties of the CVS procedure. To address the second hypothesis, we correlated effect sizes between behavioural tests.

## Materials and methods

We searched PubMed for the terms “chronic variable stress” and “chronic unpredictable stress” in the Title/Abstract field. The search was done on 31st October 2023 and included all articles published from November 2018, i.e., a five-year period. Exclusion criteria are shown in Figure 1.

**Figure 1:**
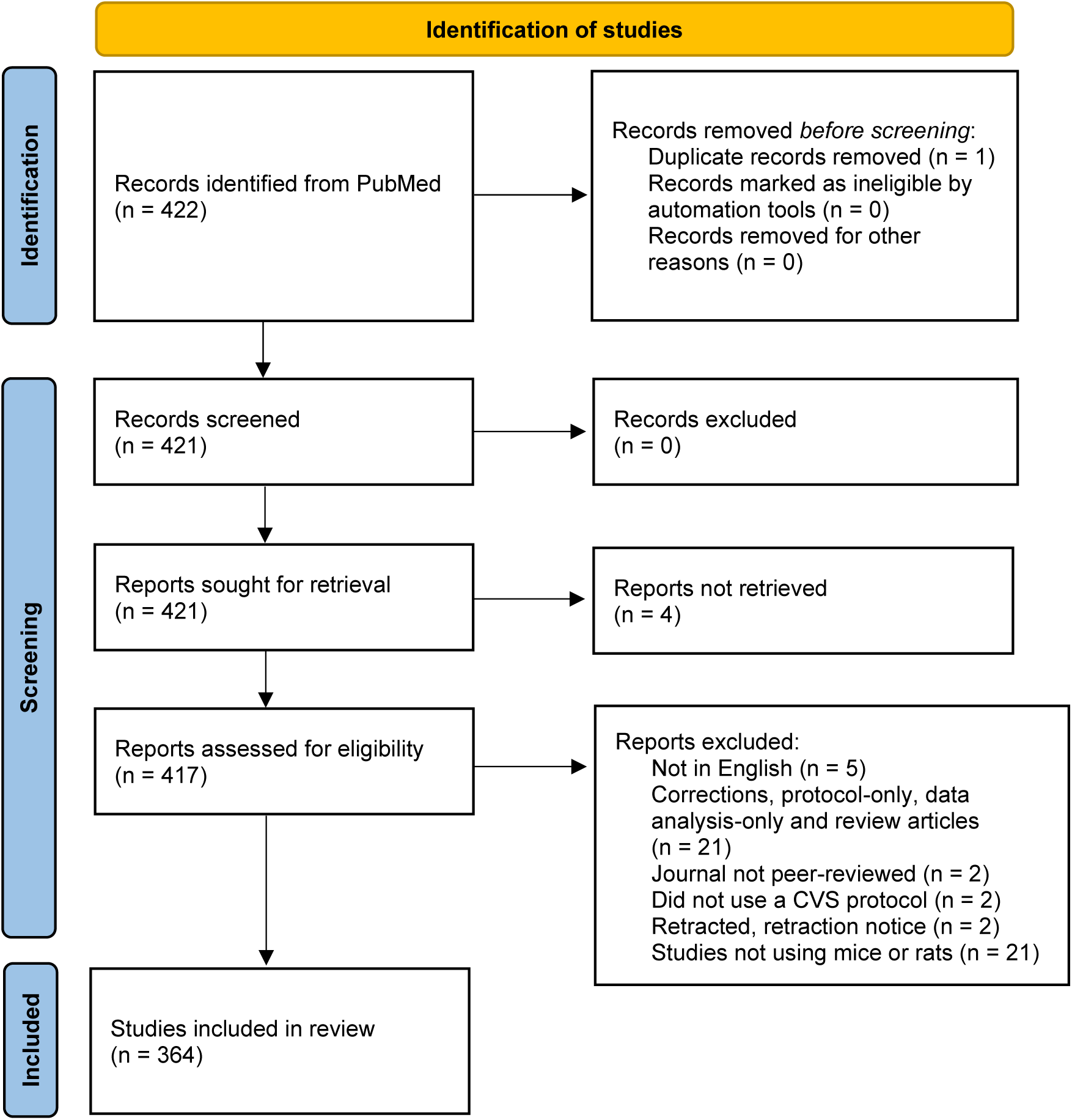
Identification of studies.

For each article using mice, we characterised the duration, burden and diversity of the CVS procedure. ‘Duration’ was the length of the CVS procedure in days. ‘Burden’ was the total number of stressors experienced by each animal during the entire CVS procedure. ‘Diversity’ was the total number of different types of stressors used. For example, if an article used five different stressors, with one of these stressors imposed once daily for a total of 10 days, we recorded a duration of ten, a burden of ten and a diversity of five. Different types of stressors may have different impacts on animals; for example, a 10 min recording of bird cries^30^ may be a much less intense stressor than a 5 min forced swim in ice-cold water^31^. However, we have no means of determining how different stimuli are perceived by other species, so we did not seek to account for potential differences in the impact of different types of stressors in our analysis. Most CVS procedures used only one particular stressor at any one time, but some used combined stressors. Combining stressors may have an effect that is greater or lesser than the sum of their parts, but for the reason given above we did not seek to account for an effect of combining stressors in our analysis. Accordingly, for articles using combined stressors, we counted each stressor separately in our calculation of the CVS burden.

We also documented whether a justification for selecting the particular CVS procedure was given in the methods section, whether that procedure had been used in an earlier article, and whether the authors had modified their CVS procedure from the one cited.

We then systematically documented the types of behavioural tests used to evaluate the effects of CVS. To understand how consistent behavioural testing was between articles, we focused on articles using mice that used the SPT, the TST, the FST, the OFT and/or the EPM and documented specific characteristics for each test. For the SPT, we recorded the concentration of sucrose used, the duration of exposure to the sucrose/water choice and whether and for how long the animals were deprived of food and/or water before the test. For the TST, we recorded the duration of the test (including any period of habituation) and the consequences of tail-climbing (i.e., escape) behaviour during the test. For the FST, OFT and EPM, we recorded the duration of the test.

In order to explore whether species differences may be relevant, we also systematically documented behavioural tests in 161 articles using CVS procedures with rats published in the same time period. We selected articles that used either the SPT and the FST or the OFT and the EPM and documented characteristics of the CVS procedures and details of the behavioural tests in the same way as described above.

To estimate the effect sizes, we used an online tool (foxyutils.com/measure-pdf) to measure the size of the mean and its error bar (representing either the standard error of the mean (SEM) or the standard deviation (SD)) in figures depicting results from these tests. If the article gave the SEM, we calculated the SD using the published sample size. If the sample size was given as a range, we used the lowest number in the range for this calculation. We used R to calculate the effect size (Cohen’s d) of CVS exposure on sucrose preference in the SPT, and on time (or proportion of time) spent immobile in the TST and FST, in the centre of the field in the OFT, or in the open arms of the EPM. Cohen’s d was calculated, as

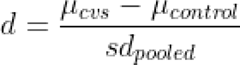

where s_pooled_ is a pooled measure of variability, calculated by:

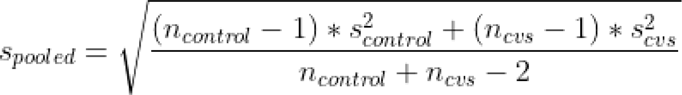

Accordingly, our calculated effect sizes have a negative sign if the mean of the CVS group is lower than the mean of the control group (for example, the typical outcome for sucrose preference in the SPT or time in the open arms in the EPM) and a positive sign if the mean of the CVS group is higher than the mean of the control group (for example, the typical outcome for time spent immobile in the FST).

We next asked whether effects sizes in each of these tests correlated with characteristics of the CVS procedure used. For each test, a linear model was generated in the form of:

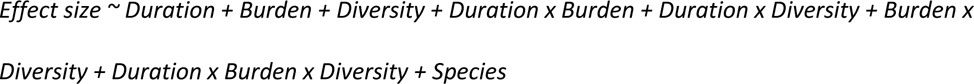

This model considers the effect of duration, burden and diversity and their possible interactions (which we considered as biologically plausible) on the effect size; we also included species as a covariate but did not explore possible interactions with the other factors. Assumptions of linear models were visually confirmed by visualization of diagnostic plots (residual vs fitted and residual QQ plots). To determine whether procedures imposing different levels of stress would result in different effect sizes, we calculated the Euclidean distance between studies’ duration/diversity/burden and correlated those with the differences in effect size between the studies.

To explore whether effect sizes correlated *between* different behavioural tests, we calculated a pairwise Pearson correlation between effect size for all possible pairs of observations. For example, if a study used FST, TST and EPM, we calculated cor(FST, TST), cor(FST, EPM) and cor(TST, EPM) for that specific study.

The code used for analysis, as well as the full dataset can be downloaded at https://github.com/nicolaromano/Romano_Menzies_2024_CVS. A reproducible analysis environment is provided through the use of the *renv* package.

We note that only five articles reported data on both males and females. We stratified effect size data by sex in those studies, but we did not systematically compare effect sizes by sex or any other characteristics of the animals. Other limitations of our study are given in Section 1 of the Supplementary Text.

## Results

We systematically documented CVS procedures used in 202 articles using mice. 192 articles (95%) gave sufficient detail to determine the duration of the CVS procedure, this ranged from 4-84 days with a median of 21 days. 153 articles (76%) gave sufficient detail to determine the burden of the CVS procedure. The median burden was 37 stressors/mouse with a range of 4-140 stressors/mouse. The median number of stressors experienced by each mouse each day was 2, with a range of 0.4-7 stressors/day. 196 articles (97%) gave sufficient detail to determine CVS diversity. The median was 8 with a range of 2-14 (Figure 2).

**Figure 2:**
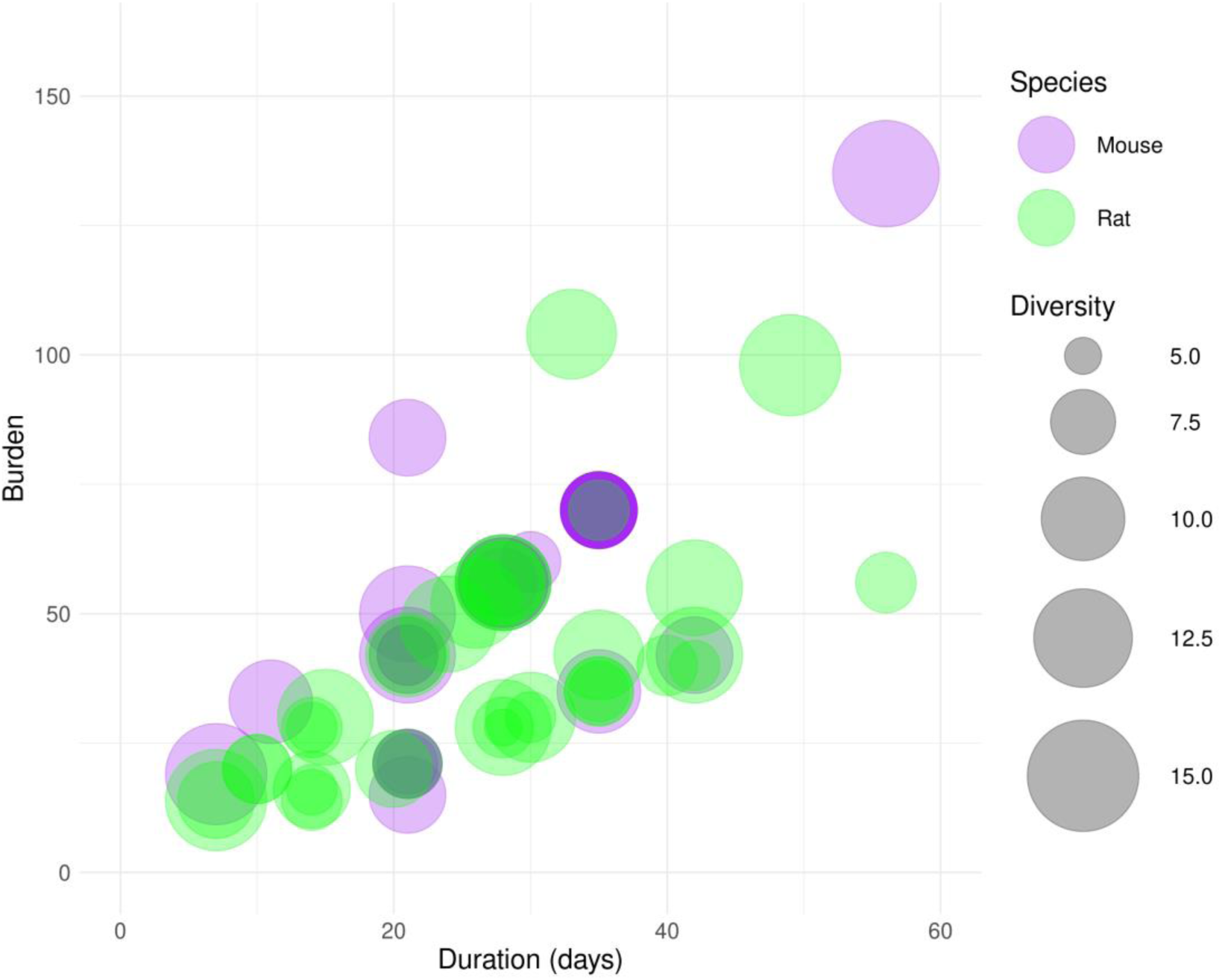
Bubble chart showing the duration, burden and diversity (bubble width) of CVS procedures used in mouse and rat studies.

Thirty-nine different stressors were used across the mouse CVS studies. The most commonly used stressors were physical restraint (used in 165 articles; 82%), changes to the light-dark cycle (142 articles; 71%), wet cage bedding (139 articles; 69%), cage tilting (138 articles; 68%), food deprivation (103 articles; 51%), water deprivation (99 articles; 49%) and a forced swim in cold water (95 articles; 47%). 164 of the 202 articles (81%) used a unique combination of stressors in their CVS procedure. Any articles that used an identical CVS procedure had authors in common, with the exception of a 21-day CVS procedure involving once-daily imposition of either restraint, foot shock or tail suspension that was used by five apparently independent groups (Supplementary Data 1).

184 articles using mice (91%) gave no justification for their choice of CVS procedure. Three articles briefly discussed the factors behind their choice of procedure^32–34^, and a further article stated that stressors were selected “to not directly perturb thermogenic, metabolic, or pain pathways”^35^. Other articles made only general comments, for example, stating that the chosen procedure prevented habituation or adaptation. 139 articles (69%) cited a previous study as a basis for their choice of CVS procedure. Fifty-four of those articles stated that the cited procedure was modified for use in their own study, but none stated what the modification was or gave detail on why it was made (Supplementary Data 1).

179 articles using mice (89%) reported using behavioural tests. A total of thirty-seven different post-CVS behavioural tests were reported across these articles, with a mode of three tests per article (range 1-8). The most commonly used tests were the FST (used in 122 articles; 60%), the OFT (93 articles; 46%), the SPT (84 articles; 41%), the TST (79 articles; 37%), and the EPM (49 articles; 26%). All other behavioural tests were used in twenty-nine or fewer articles (Supplementary Data 2).

To explore correlations between effect sizes, we pooled effect size data from all mouse studies that used either (1) the SPT, the TST and the FST, or (2) the OFT, the EPM and the FST in that particular study. First, we asked whether these tests were done in a consistent way. We found remarkable variability in how tests were done, noting nineteen unique variations in how an SPT was performed, ten unique variations in how a TST was performed and five unique variations in how an FST was performed. There were three unique variations in how an OFT and an EPM were performed, but differences were based solely on the duration of the test (Supplementary Tables 1-5). Broadly speaking, exposure to CVS was associated with a reduction in sucrose preference, an increase in the time spent immobile in the TST and FST, and a reduction in the time spent in open areas in the OFT and EPM. However, one study^36^ reported a small increase in the mean sucrose preference in CVS-exposed mice, one^37^ reported a small reduction in immobility in the TST in CVS-exposed mice, and one^33^ reported an increase in time spent in the open arms of the EPM in CVS-exposed mice. A large range of effect sizes was seen. The median effect size (Cohen’s d) in the SPT was −1.70 (range: −3.98;0.56), 1.46 (range: −0.28;4.18) in the TST, 1.74 (range: −0.53;5.96) in the FST, −0.96 (range: −3.84;-0.06) in the OFT, and −0.95 (range: −4.57;0.55) in the EPM (Supplementary Data 2).

108 articles using rats (67%) reported using behavioural tests. A total of thirty different behavioural tests were reported. The mode was three tests per article (range 1-6). The most commonly used tests were the OFT (used in 59 articles; 37%), the SPT (58 articles; 36%), the FST (52 articles; 32%), and the EPM (38 articles; 24%). All other behavioural tests were used in thirteen or fewer articles (Supplementary Data 4). Only five rat studies (3%) used the TST, so to explore corelations between effect sizes, we focused on studies that used the SPT and the FST, or the OFT and the EPM. We documented twenty-two unique variations in how an SPT was performed and six unique variations in how an FST was performed. There were just two and three unique variations respectively in how an OFT and an EPM was performed, with differences based solely on the duration of the test (Supplementary Tables 6-9).

As with mouse studies, a large range of effect sizes was seen in the behavioural tests used to evaluate the effects of CVS in rats. The direction of change in the tested variable was as expected, with the exception of two studies where the time spent in the open arm of the EPM was higher in the CVS group compared to the control group^38, 39^, and one study where the time spent in the central area of the OFT was higher in the CVS group compared to the control group^40^. The median effect size in the SPT was −1.80 (range −7.61;-0.18), 1.59 (range 0.08;6.55) in the FST, −0.76 (range −4.55;2.42) in the OFT, and −0.88 (range −3.17;0.87) in the EPM (Supplementary Data 5).

We characterised the CVS procedures in the subset of fifty-four rat studies that used the SPT and the FST, or the OFT and the EPM (Figure 2). We did not systematically record all stressors used in these rat studies, but we noted that these articles used similar stressors to those documented for mouse studies. One exception was an article that modelled Gulf War Illness by giving oral pyridostigmine and intranasal lipopolysaccharide 5-7 times per week alongside a 33-day CVS procedure^41^. Fifty-two articles (96%) gave sufficient detail to determine the duration of the CVS procedure, this ranged from 7-63 days with a median of 28 days. Forty-three articles (80%) gave sufficient detail to determine the burden of the CVS procedure, the median was 38 stressors/rat with a range of 10-126 stressors/rat. Fifty-two articles (96%) gave sufficient detail to determine diversity. The median diversity was 10 with a range of 5-15. All but six of these fifty-four articles used a unique CVS procedure (Supplementary Data 5).

We took advantage of the lack of consistency in CVS procedures used to ask whether effect sizes in behavioural tests were related to the duration, burden and diversity of CVS procedures (Figure 3). Adjusted R^2^ values indicated that no more than 62% of the variability in effect sizes was explained by the duration, burden and diversity of the CVS procedures in any behavioural test (Tables 1-5). The “Estimate” value in Tables 1-5 is an estimate of the slope of the relationship between one or more CVS characteristics and the effect size in that test. A summary of this analysis is given in Figure 4. We found statistically significant interactions between the three variables describing CVS characteristics, but these were not consistent between tests (Tables 1-5). We interpreted interactions as complex relationships between variables; for example, in the SPT, there was a significant interaction between duration and burden. This means that the SPT effect size depends on the CVS duration, but in a way that is affected by CVS burden. Because of the interaction, we ignored the coefficient for either single term (Duration or Burden alone). None of the three CVS characteristics, or their interactions, had a significant effect on effect size magnitude in the TST or OFT. There were no significant differences with species as a factor for any behavioural test (“Species (rat)” in Tables 1-5).

**Figure 3:**
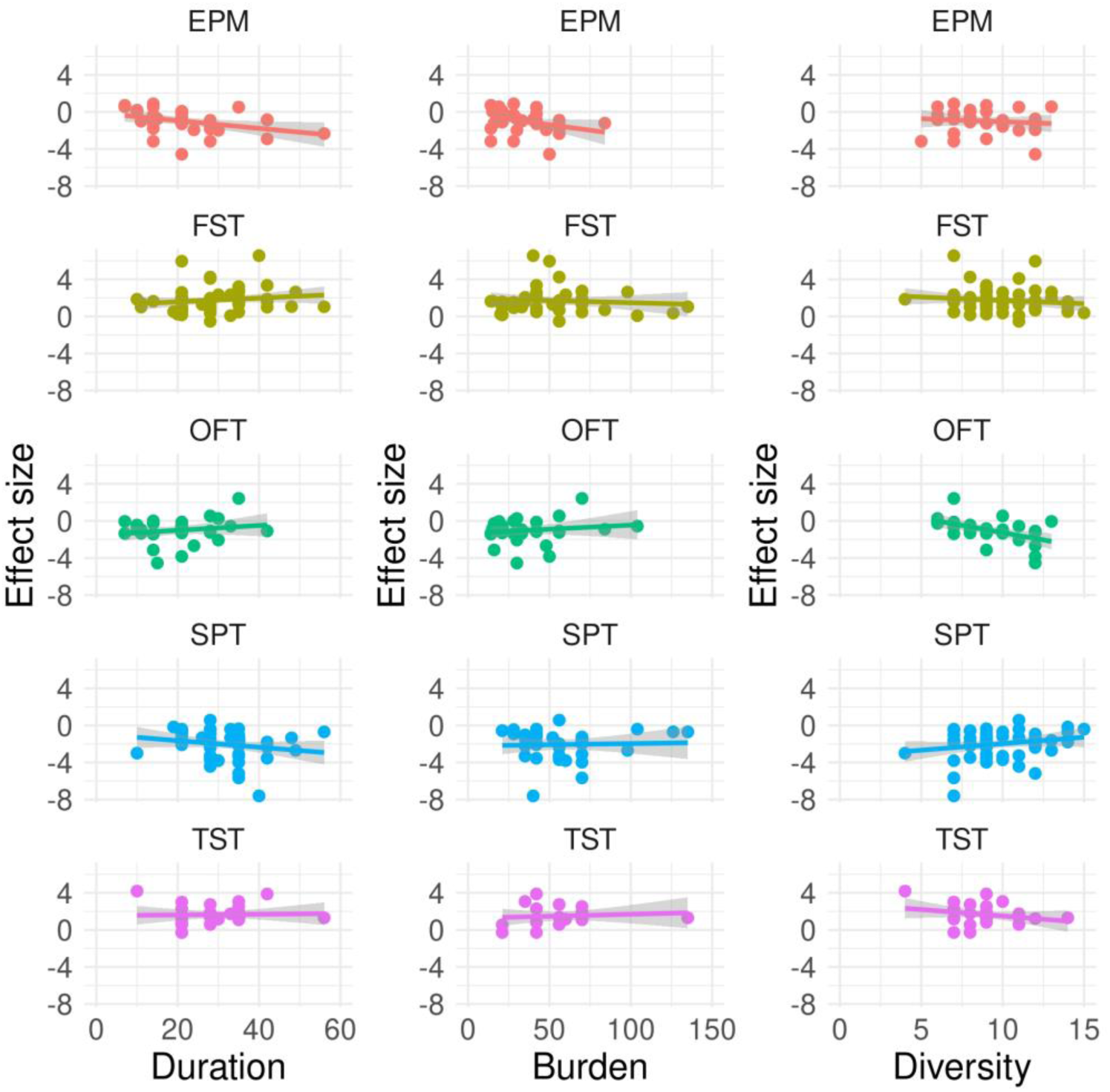
Correlation between effect sizes in the SPT, TST and FST, and OFT, EPM and FST. Each data point represents an article. A linear fit is shown. The grey band indicates 95% confidence intervals.

**Figure 4:**
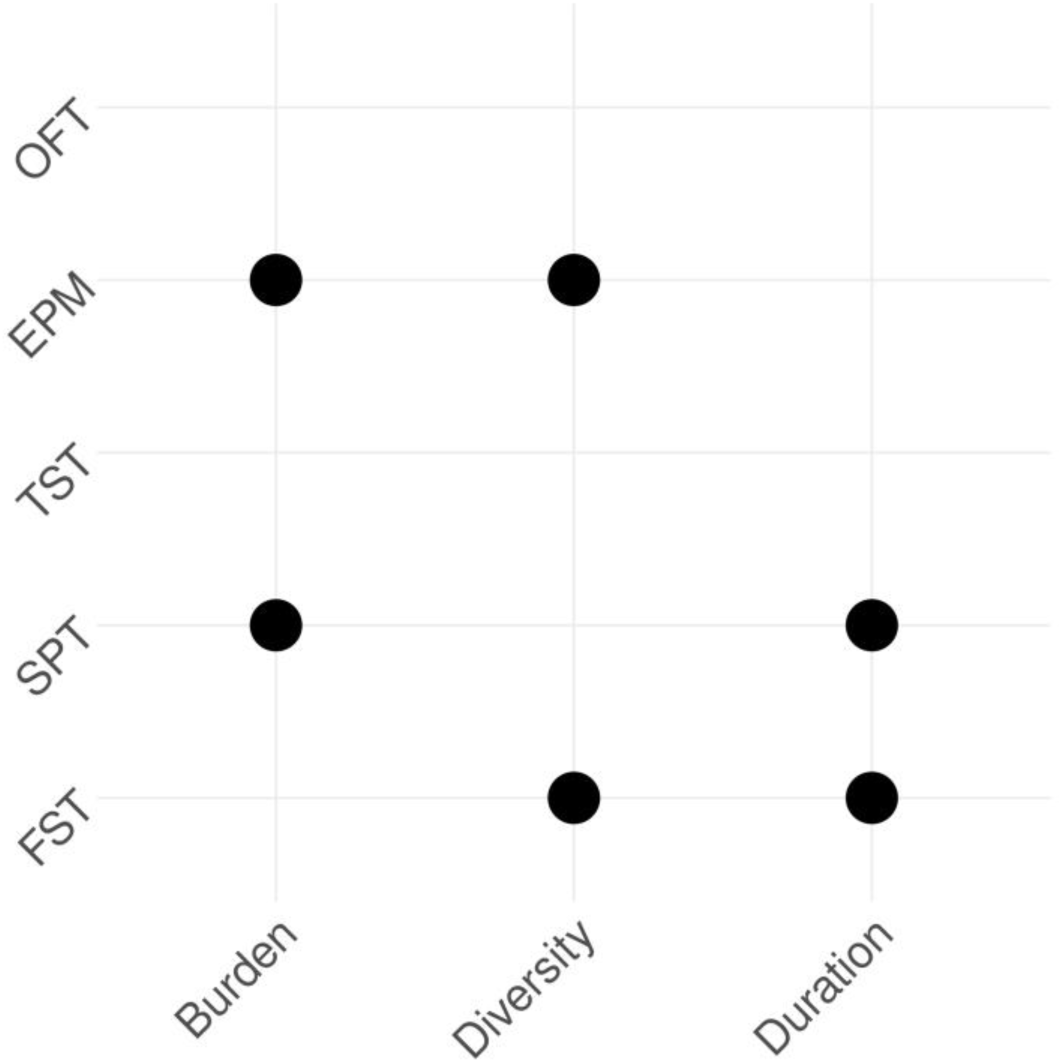
Summary of CVS characteristics’ influence on effect sizes. A dot is shown where a significant interaction between CVS characteristics was detected for that test.

**Table 1:** Estimates for effect sizes in the sucrose preference test. Adjusted R^2^ = 0.412.

|  | Estimate | SEM | t value | p value |
| --- | --- | --- | --- | --- |
| (Intercept) | 20.126 | 12.557 | 1.603 | 0.121 |
| Duration | -0.536 | 0.396 | -1.354 | 0.187 |
| Burden | **-0.484** | **0.214** | **-2.265** | **0.032** |
| Diversity | -1.635 | 1.430 | -1.143 | 0.263 |
| Species (rat) | -0.634 | 0.571 | -1.109 | 0.277 |
| Duration x Burden | **0.010** | **0.005** | **2.257** | **0.032** |
| Duration x Diversity | 0.035 | 0.046 | 0.761 | 0.453 |
| Burden x Diversity | 0.043 | 0.022 | 1.941 | 0.063 |
| Duration x Burden x Diversity | -0.0008 | 0.0005 | -1.772 | 0.088 |

**Table 2:**
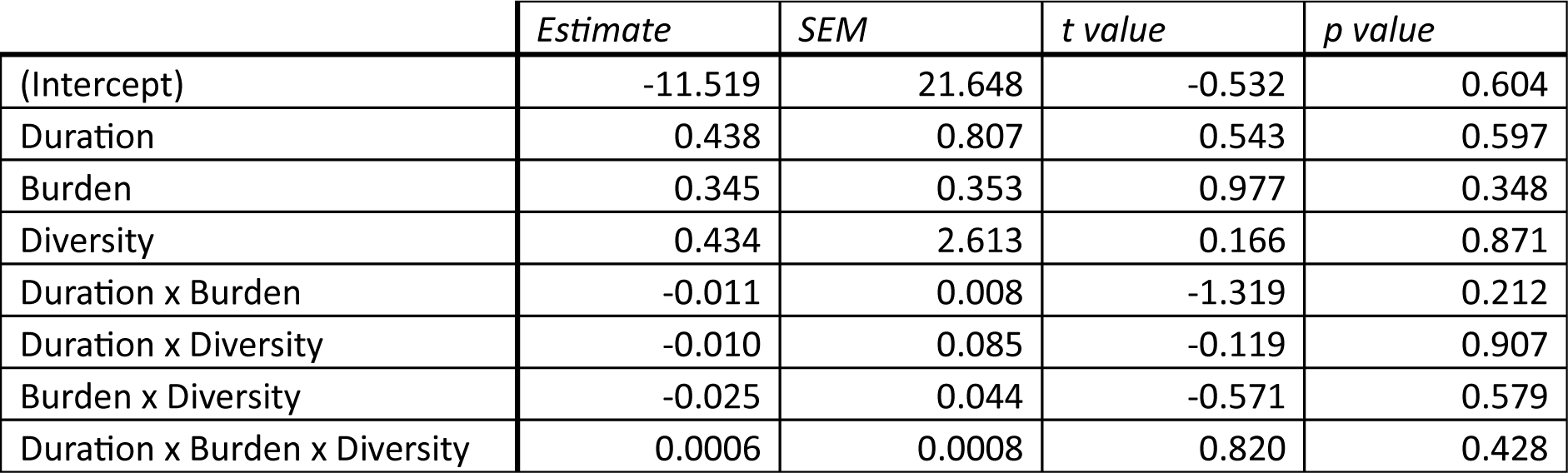
Estimates for effect sizes in the tail suspension test. Adjusted R^2^ = 0.652. These values are based on data from mouse studies only.

|  | Estimate | SEM | t value | p value |
| --- | --- | --- | --- | --- |
| (Intercept) | -11.519 | 21.648 | -0.532 | 0.604 |
| Duration | 0.438 | 0.807 | 0.543 | 0.597 |
| Burden | 0.345 | 0.353 | 0.977 | 0.348 |
| Diversity | 0.434 | 2.613 | 0.166 | 0.871 |
| Duration x Burden | -0.011 | 0.008 | -1.319 | 0.212 |
| Duration x Diversity | -0.010 | 0.085 | -0.119 | 0.907 |
| Burden x Diversity | -0.025 | 0.044 | -0.571 | 0.579 |
| Duration x Burden x Diversity | 0.0006 | 0.0008 | 0.820 | 0.428 |

**Table 3:** Estimates for effect sizes in the forced swim test. Adjusted R^2^ = 0.290.

|  | Estimate | SEM | t value | p value |
| --- | --- | --- | --- | --- |
| (Intercept) | -18.281 | 9.104 | -2.008 | 0.052 |
| Duration | **0.787** | **0.264** | **2.976** | **0.005** |
| Burden | 0.113 | 0.173 | 0.655 | 0.517 |
| Diversity | 1.912 | 0.974 | 1.963 | 0.058 |
| Species (rat) | -0.499 | 0.436 | -1.144 | 0.261 |
| Duration x Burden | -0.006 | 0.004 | -1.577 | 0.124 |
| Duration x Diversity | **-0.073** | **0.028** | **-2.617** | **0.013** |
| Burden x Diversity | -0.011 | 0.018 | -0.591 | 0.558 |
| Duration x Burden x Diversity | 0.0005 | 0.0004 | 1.400 | 0.170 |

**Table 4:** Estimates for effect sizes in the open field test. Adjusted R^2^ = 0.623.

|  | *Estimate* | *SEM* | *t value* | *p value* |
| --- | --- | --- | --- | --- |
| (Intercept) | -1.567 | 5.607 | -0.280 | 0.782 |
| Duration | -0.025 | 0.241 | -0.103 | 0.919 |
| Burden | 0.150 | 0.193 | 0.779 | 0.444 |
| Diversity | 0.3034 | 0.564 | 0.538 | 0.596 |
| Species (rat) | -0.704 | 0.428 | -1.643 | 0.115 |
| Duration x Burden | 0.00007 | 0.007 | 0.011 | 0.991 |
| Duration x Diversity | -0.008 | 0.024 | -0.328 | 0.746 |
| Burden x Diversity | -0.025 | 0.020 | -1.275 | 0.216 |
| Duration x Burden x Diversity | 0.0004 | 0.0007 | 0.612 | 0.547 |

**Table 5:** Estimates for effect sizes in the elevated plus maze. Adjusted R^2^ = 0.482.

|  | *Estimate* | *SEM* | *t value* | *p value* |
| --- | --- | --- | --- | --- |
| (Intercept) | -7.294 | 5.262 | -1.386 | 0.179 |
| Duration | -0.035 | 0.266 | -0.133 | 0.895 |
| Burden | **0.422** | **0.164** | **2.566** | **0.017** |
| Diversity | 1.031 | 0.585 | 1.762 | 0.091 |
| Species (rat) | -0.192 | 0.438 | -0.438 | 0.665 |
| Duration x Burden | -0.007 | 0.007 | -1.093 | 0.286 |
| Duration x Diversity | -0.009 | 0.031 | -0.282 | 0.780 |
| Burden x Diversity | **-0.053** | **0.019** | **-2.832** | **0.009** |
| Duration x Burden x Diversity | 0.001 | 0.0008 | 1.342 | 0.193 |

***Tables 1-5:*** *Estimations of the effects of species, CVS duration, burden and diversity on the magnitude of effect size in five behavioural tests commonly used to evaluate the effects of CVS; p<0.05 is shown in bold*.

To explore how differences in CVS procedure affect effect sizes, we calculated how distant studies were from each other in terms of duration, burden and diversity of their CVS procedure. We reasoned that if these aspects of procedure design related to the impact of CVS, then a large distance between procedure designs would be associated with differences in effect sizes. Figure 5 shows the relationship between distance in procedure design and difference in effect size for the FST. In the majority of cases, a larger difference in procedure design did not correspond to a larger difference in effect size. Overall, there was no significant correlation between difference in procedure design and difference in effect size (correlation = 0.01, p = 0.668, correlation test; see Section 2 of the Supplementary Text for specific examples).

**Figure 5:**
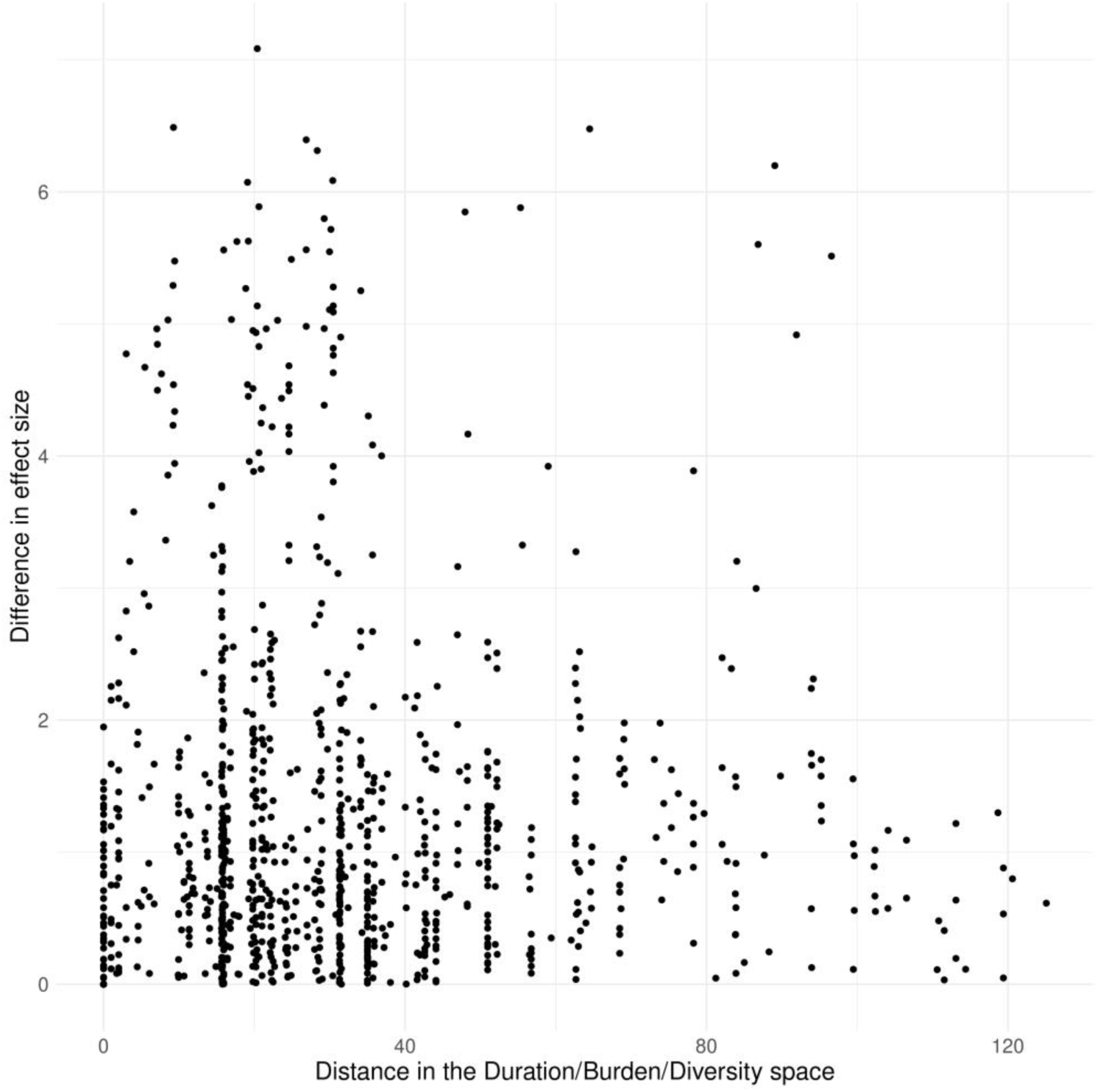
Correlation between the Euclidean distance of CVS characteristics and the difference in FST effect size for all study pairs.

To explore whether evaluative behavioural tests were measuring the same or different consequences of CVS, we next asked whether effect sizes correlated between tests (Figure 6). The TST and FST both measure immobility, and we found a moderate positive correlation in effect sizes between these two tests (Pearson’s correlation coefficient (r) = 0.57). The OFT and EPM are both deemed to measure anxiety, and we found a strong positive correlation in effect sizes between these two tests (r = 0.74). There was a very strong positive correlation in effect sizes between the EPM and FST (r = 0.91) and between the EPM and TST (r = 0.97). There was also a very strong positive correlation in effect sizes between the OFT and FST (r = 0.94), but not between the OFT and TST (r = −0.12). There was a moderate positive correlation between SPT effect sizes and TST, FST and EPM effect sizes (r = 0.62, 0.64 and 0.45 respectively), but no correlation with OFT effect size (r = −0.01).

**Figure 6:**
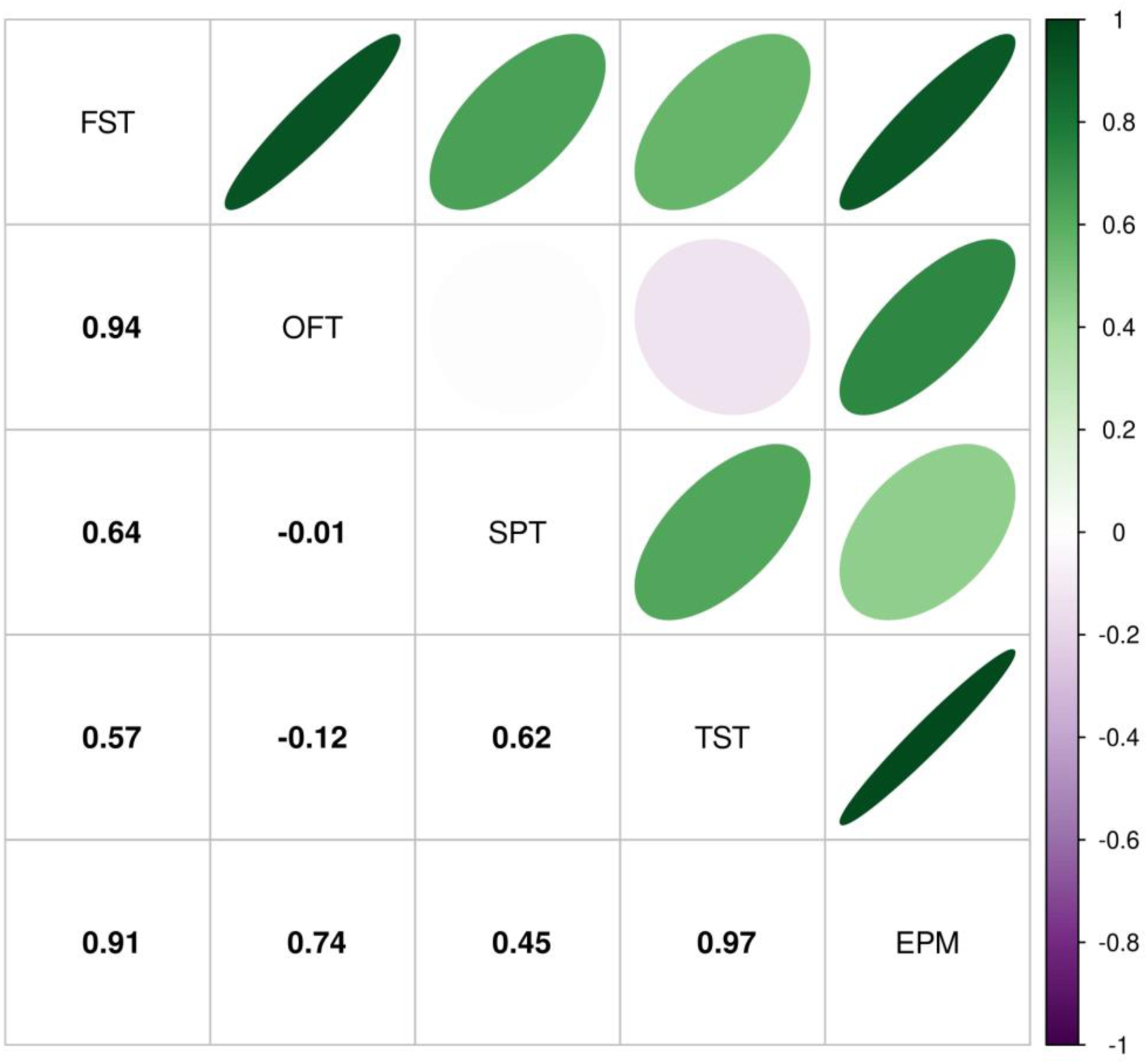
Correlations (Pearson’s correlation co-efficient) between SPT, TST, FST, OFT and EPM effect sizes.

## Discussion

CVS procedures deliberately impose suffering. Because of this, researchers should take particular care with study design. However, our study revealed a considerable and largely unexplained variability in the design of CVS procedures. In more than half of the 202 mouse studies included in our study, each animal experienced more than fifty separate episodes of stress. In more than a quarter, stressors were regularly imposed for more than a month. In most studies, stressors were imposed at least once every day, and in one case, up to four times per day^42^. Most authors did not explicitly justify their choice of CVS procedure, regardless of whether they used or modified a previously published procedure. However, we noted one article using rats that stated that stressors were chosen because they had been previously used, they were known to be “mild” but have “well-known harmful effects”, and they were feasible to carry out^40^. Similar reasons for stressor selection were given in an article describing a CVS study design for flies^43^. As such, it seems little scientific justification is normally given for the specific design of a CVS procedure, including how the design might be relevant to stress-related conditions in humans. But does this variability in procedure design matter? Perhaps researchers just want to impose stress, and the detail of the procedure is unimportant?

To be clear, we do not endorse an arbitrary approach to study design, and others have noted the relevancy of CVS duration (suggesting a maximum duration of ten days)^44^, the optimal hiatus between the final stressor and the evaluation of the procedure (ideally ∼24 hours)^44^, potential confounds of single-versus group-housing^44^, the lack of studies on female animals^26, 44^, the focus on short-term effects when these procedures aim to model a long-term condition^45^, and how ‘normal’ control animals actually are^46, 47^. However, apart from classifying animals as “stress-susceptible” or “stress-resilient” according to behaviours in evaluative tests^27, 48^, there has been relatively little investigation into whether the number, type, order, pattern or frequency of stressors differentially affect performance in common evaluative behavioural tests^44, 49–51^. Similarly, very few studies determine a time course for the effects of CVS, potentially because many of the evaluative tests are themselves stressors. However, a few studies have carried out multiple SPTs during a CVS procedure, and qualitative variation is reported. For example, a step-like decrease in preference after 28 days^52^, or a gradual ramping down in preference over nine weeks that gradually and partially reverses over five weeks after CVS offset^53^. In a systematic study^54^, sucrose preference was measured weekly during a nine-week CVS procedure in three strains of mice. Differences in preference were seen between control and stressed mice only in weeks 2, 5 and 9 in CBA/H mice (though in week 9 the stressed mice had a higher sucrose preference than controls), and in week 5 in the C57BL/6 and the DBA/2 mice (the stressed C57BL/6 mice had a higher sucrose preference than controls). As such, there seems to be little understanding of how CVS duration relates to sucrose preference behaviour.

We also documented a lack of consistency in tests used to evaluate the effects of CVS. Across 202 mouse articles, thirty-seven different post-CVS behavioural tests were used, and two or more of the five most commonly-used tests were used in twenty-two unique combinations. In addition, there was a lack of consistency in the design of a given behavioural test. Taking the SPT as an example: efforts have been made to standardise this popular and ostensibly simple test^26, 55^, but in just twenty-seven articles using mice we noted eighteen unique variations in how an SPT was done, with the test duration ranging from one to forty-eight hours, and one of eight different patterns of food and/or water deprivation being imposed prior to the test (Supplementary Table 1). Another report has described over 100 variations in how the SPT is done^25^. Variation in the duration of immobility tests may also be important. If at some point in the test the animal chooses to become and remain immobile, the longer an immobility test is, the larger the effect size will be. For example, one study^56^ reported an FST where control mice were immobile for ∼70 seconds and CVS-exposed mice were immobile for ∼900 seconds; we estimated the effect size to be 21.2. In order to be able to directly compare FST effect sizes, clearly all tests need to be of the same duration, and the time spent mobile and the latency to immobility should also be reported. If immobility occurs in bouts, this should be carefully quantified and reported.

Setting these issues aside, we wanted to explore whether there was a relationship between CVS characteristics and the magnitude of effect sizes measured in commonly-used behavioural tests. None of the five behavioural tests were sensitive to all three CVS characteristics. We observed very few significant correlations between CVS characteristics (or interactions between two or more characteristics) and effect sizes. Those that were significant were small in magnitude (<0.1 ‘units’ of Cohen’s d), and for the FST and EPM, they were negative. In other words, increasing the duration or intensity of a CVS procedure has only a small effect on effect sizes, and for two tests, increasing the duration or intensity of a CVS procedure is associated with decreases in effect size. Our analysis indicates that effect sizes in only one of five commonly-used behavioural tests – the FST – is sensitive to variations in CVS procedure design, albeit in a complex way. However, when exploring how variation in the CVS procedure affected FST effect size, we did not find a clear correlation between the difference in effect size and the distance between the procedures.

This disconnect may be because different CVS procedures *are* able to influence effect size magnitude, but we did not detect those correlations because the CVS characteristics we quantified are not the characteristics that influence effect size. For example, the OFT was the only test where we observed a relatively large R^2^ value. However, we could not detect any significant effects of CVS procedure characteristics on OFT effect size. This could be because (1) there are other factors influencing effect size that we did not include in our analysis, or (2) the factors we chose *are* important, but a linear model is not an appropriate way of detecting them (though inspection of the data does not suggest this). With respect to (1), perhaps the *type* of stressor used is the only key factor. In other words, certain types of stressors evoke large effect sizes, and other types evoke small effect sizes. This could be readily tested by fixing duration and burden and comparing different stressors, potentially using existing data from chronic restraint stress studies^57^. Alternatively, it is possible that differences in diversity, burden and duration *do* evoke different behavioural states, but these states (or differences in the depths of these states) are not readily detected by commonly-used behavioural tests.

We next asked whether effect sizes correlated between tests. This was partly motivated by uncertainty around what these behavioural tests actually measure^21^. For example, a recent systematic review^26^ noted that the availability of food and water during an SPT is essential in order to be confident that sucrose preference is related to stress-sensitive reward rather than hunger or thirst (we note here that the majority of mouse and rat studies we documented withheld food and/or water prior to the SPT). However, the question of what an SPT measures, even in non-deprived conditions, is still open. Some authors have concluded a reduction in sucrose preference is related to stress-induced changes in learning and memory rather than anhedonia^58, 59^. Importantly, sucrose preference data is often reported without key information on the volume of water and sucrose solution consumed. For example, a detailed analysis of sucrose/water choice in mice after CVS reported a reduction in sucrose preference from ∼85% to ∼68%^59^. However, total fluid consumption increased from ∼3.3 ml to ∼4.1 ml, water intake increased from ∼0.5 ml to ∼1.3 ml, and sucrose intake was stable (∼2.8 ml), meaning sucrose preference decreased solely because water intake increased.

We found that effect sizes in the TST and FST (tests that both measure immobility) correlated positively, albeit only moderately. The most parsimonious explanation is that the post-CVS changes that underlie the immobility measured in one test also underlie the immobility measured in the other test. However, we saw that FST effect sizes are sensitive to CVS duration and diversity, but TST effect sizes are not, suggesting some divergence in what these tests measure and/or the sensitivity with which they measure it. Effect sizes in the OFT and EPM (tests that are both believed to measure anxiety) showed a strong positive correlation, again suggesting that these tests measure the same thing. However, EPM effect sizes correlated much more strongly with TST and FST effect sizes than with OFT effect sizes, and a meta-analysis of mouse behavioural tests showed that measuring time spent in the open arms of the EPM, but not time spent in the centre of the OFT, was an effective assay for an anxiolytic effect^24^, suggesting that there may be some divergence in what the EPM and OFT measure. One simple interpretation of the strong EPM-TST-FST correlation is that the EPM measures TST/FST-sensitive consequences of CVS more strongly than OFT- sensitive consequences.

We found a strong positive correlation between SPT effect size and TST effect size, and between SPT effect size and FST effect size. This suggests that the SPT also measures the TST/FST-sensitive consequences of CVS. In line with this, there was a modest correlation between SPT and EPM effect sizes, though the absence of a correlation between SPT and OFT effect size complicates the picture.

Variability in CVS procedure design and evaluation matters ethically as well as scientifically. Without evidence for procedure-dependent effects on behaviour, a researcher may choose (incorrectly in our view) to impose a long and severe CVS procedure in the hope it will cause a larger effect size, but the use of long and severe procedures raises ethical concerns^60^. This is crystallised in a statement made in an article describing a statistically non-significant finding as something that could “potentially be mitigated by augmenting the sample size and duration of stress protocol”^61^. Flagging this comment is not intended as a barb against these authors in particular; similar comments on sample size are widespread in the biomedical literature. For example, we recently carried out a systematic review (unpublished) to explore one of the canonical Three Rs: reduction. We documented how sample size was determined in 385 translational CVS studies (including all the rodent articles included here). We found that 92% of articles did not report how the sample size was determined, and that only one article used a biologically meaningful effect size to calculate sample size. Ignoring the principle of reduction in order to achieve ‘statistical significance’ is problematic from both an ethical and a scientific standpoint; arbitrarily increasing sample size just in order to detect a small, possibly biologically irrelevant, effect size is not advisable^62^.

Researchers who use CVS procedures presumably do this work because they believe these studies have translational potential. That belief surely entails accepting that the animals used in these procedures suffer physically and mentally in the same or similar way to humans who are exposed to uncertainty, fear, pain, hunger or thirst^63^ (though with the exception of the article that set out to model Gulf War Illness^41^, none of the articles clearly stated what human experience they were modelling). If researchers do believe animals suffer during CVS procedures, these studies must be designed so their benefits outweigh the harms. This can only be done if we have some confidence in what the evaluative tests measure, and if we use stressors that we know can influence the behaviours tested. None of that necessarily guarantees translatability to humans, of course^3, 28, 44^. Currently, it seems we have only partial evidence for what behavioural tests measure and which stressors modify those measures, and whatever the explanation for the apparent disjunction between stress entity and behavioural impact, our data strongly suggests that standardisation of CVS study design is needed in order for this work to be useful.

The initial step would be agreement on a standardised CVS duration and burden (say a ten-day duration^44^ with one stressor imposed per day) and the specific stressors used. We do not propose here what types of stressors should be used, or in what order, but there should be sufficient variation to avoid loss of external validity^64^, and ethically it seems most reasonable to first test stressors that activate the physiological stress response without causing significant pain or suffering. The next step would be to select an appropriate and standardised behavioural test(s) to evaluate the behavioural effect of those stressors. But what should this test be?

Taken together, our data suggest that the FST, TST, SPT and EPM all measure the same or similar consequence of CVS. This raises the question of whether animals should be required to participate in all of these tests. We argue that they should not, particularly if participation involves suffering. We note that around half of the mouse studies in our review used the FST as a stressor during the CVS procedure, and around a quarter used the TST in the same way, implying that many researchers believe these tests do cause suffering. There are doubts around the FST’s face, construct, internal, external and predictive validity as a measure of stress-induced depression^21, 65–69^. In 2023, the UK Government recommended that the FST “should not be used as a model of depression or to study depression-like behaviour […] or for studies of anxiety disorders and their treatment”^70^, and there are plans to ban the FST in the UK^71^. Our effect size correlations indicate the FST measures the same (unidentified) thing(s) as the TST and EPM (though see Figure 4; none of these three tests seem to be sensitive to the same set of CVS characteristics). However, we also show that SPT effect sizes correlate well with the EPM, TST and FST, so the SPT may the best choice of test.

In contrast to the EPM and OFT, which both require specialist apparatus and recording/tracking equipment, performing and quantifying the SPT only needs bottles and a balance. Also in contrast to the TST and FST, carrying out an SPT is unlikely to induce an acute stress response and potentially confound the wider study, at least in sucrose-habituated animals. Importantly, our analysis suggests SPT effect size is sensitive, albeit only in a very slight way, to CVS duration and burden, but insensitive to diversity. This means that the unpredictable element of an optimised CVS procedure should be the timing and duration of individual episodes of stress, rather than the supposed unpredictability arising from using a wide variety of different types of stressors (particularly given that stressors are currently often imposed at regular, therefore predictable, times of day). Using fewer, and milder, types of stressor could improve CVS procedures in terms of the amount and cost of equipment needed to impose stress, and lessen the psychological impact on researchers and carers of imposing severe stressors^72^.

Accordingly, we recommend that researchers focus on using the SPT to evaluate CVS procedures. We support the recent recommendation^26^ to use a two-bottle choice paradigm over 12 hours during the dark phase of the light-dark cycle. Rodents are most active in the dark phase, and a longer exposure period will help minimise the impact of any individual-level differences in appetitive behaviours and patterns of intake.

In addition, we recommend that a fuller data set is reported for all SPTs: not just the % preference for sucrose, but also the amount of water (ml) and sucrose solution (ml and kJ) consumed, the amount (kJ) of food consumed during the two-bottle exposure, as well as regular measurements of the animals’ 24-hour food intake and body weights. To avoid stress associated with single-housing, the experimental unit could be a pair or small group of animals housed together rather than as individuals. If studies are designed to induce and detect a biologically meaningful effect size (for example, Cohen’s d = 2.0; see Section 3 of the Supplementary Text for an explanation of why we chose this value), the required sample size could be as small as 2-3 cages per group, easing Three Rs-related concerns around using more than one animal as the experimental unit. Collecting data repeatedly over time, in the home cage, in a within-subjects design, and using a mixed-effects analysis^73^ will give a more complete picture of behaviour and energy balance during the test, allowing researchers to make better-informed inferences about the animals’ motivations for their behaviours prior to, during and after exposure to CVS.

Here we report large variability in the ways CVS studies are done – both in the design of the CVS component and in the design of the evaluative behavioural tests. Most of the studies in our review sought evidence for interventions that would prevent or reverse the effects of chronic stress. But if we are to have any confidence that translational CVS studies provide a foundation for potential clinical interventions, we must take an evidence- and ethics-informed approach to their design.

## Supporting information

Supplementary text - Romano_Menzies

Supplementary Data 01 - CVS procedures used in mouse studies

Supplementary Data 02 - Behavioural tests used in mouse CVS studies

Supplementary Data 03 - Mouse - Effect sizes in the SPT, TST, FST, OFT and EPM

Supplementary Data 04 - Behavioural tests used in rat CVS studies

Supplementary Data 05 - Rat - CVS characteristics and effect sizes in the SPT, FST, OFT and EPM

Supplementary Tables 1-9 - SPT, TST, FST, OFT and EPM characteristics in mouse and rat studies

## Acknowledgements

This study was not supported by external funding.

## Description of Supplementary Data

**Supplementary Data 1**: Characteristics of mouse CVS procedures. A light grey cell indicates the article in that row used the stressor in that column. Dark grey cells indicate that the article did not provide this information. Articles using more than one study population or experimental approach are highlighted in blue. Articles that used an identical CVS procedure are highlighted in the same colour. Data on justification of CVS procedure choice are given in columns AY-BA.

**Supplementary Data 2**: Behavioural tests used to evaluate mouse CVS studies. A light grey cell indicates the article in that row used the behavioural test in that column. A dark grey row indicates that no behavioural tests were used to evaluate the effects of the CVS procedure in that article.

**Supplementary Data 3**: Effect sizes in mouse behavioural tests. Articles using more than one study population or experimental approach are highlighted in blue.

**Supplementary Data 4**: Behavioural tests used to evaluate rat CVS studies. A light grey cell indicates the article in that row used the behavioural test in that column. A dark grey row indicates that no behavioural tests were used to evaluate the effects of the CVS procedure in that article.

**Supplementary Data 5**: CVS characteristics and behavioural test effect sizes in rat studies. Articles using more than one study population or experimental approach are highlighted in blue. Articles that used an identical CVS procedure are highlighted in the same colour.

**Supplementary Tables 1-9**: Characteristics of the SPT, TST, FST, OFT and EPM used in mouse and rat studies.

