## Supplementary text - Romano_Menzies for "Rodent chronic variable stress procedures: a disjunction between stress entity and impact on behaviour"

**Romanò & Menzies**

**Supplementary Text**

### *1. Limitations*

Our study has a number of limitations.

1. We did not preregister our protocol design[1].
2. Strict inclusion criteria may not capture all relevant articles. Setting these criteria is intended to reduce selection bias, but translational procedures that impose stressors over long periods are known by a variety of names[2], not just by our search terms, so the included articles represent a sample of all relevant studies published in that time frame. For example, we did not capture two studies cited in the main text when discussing behavioural tests[3,4], even though they fell into our time range and used and evaluated a CVS protocol in mice. This was because the title or abstract did not contain the search terms "chronic variable stress" or "chronic unpredictable stress".
3. Study inclusion/exclusion and data collection for effect size calculation was done manually by a single researcher. This may introduce biases[5,6].
4. Others have noted that the CVS literature tends to focus on adult males despite the fact that depression is more common in women than men[7], and that many of the effects of CVS seen in male rodents are not seen in female rodents[8], but we did not carry out a systematic study of the impact of sex, age, strain or any other animal characteristics on effect sizes.

### *2. Euclidean distance and effect sizes: examples of disjunction*

Here we highlight three examples from the data shown in Figure 5 to illustrate how differences in CVS procedures do not relate to differences in FST effect sizes. The first example (marked as A in Figure B below) shows two studies[9,10] with comparable CVS procedures that, despite having the same duration, burden and a similar diversity (11 vs 8) show very different effect sizes (-0.53 and 4.2). The second example (marked as B) shows how a study using a mild CVS procedure[11] (duration 40, burden 40, diversity 7) resulted in a larger effect size than one with a more intense CVS procedure[12] (duration 63, burden 126, diversity 10). The third example (marked as C) shows two studies using different procedures resulting in similar effect sizes (1.65 vs 1). In this case, the milder procedure[13] had duration 21, burden 15 and diversity 9, and the more intense procedure[14] with duration 56, burden 135 and diversity 14.

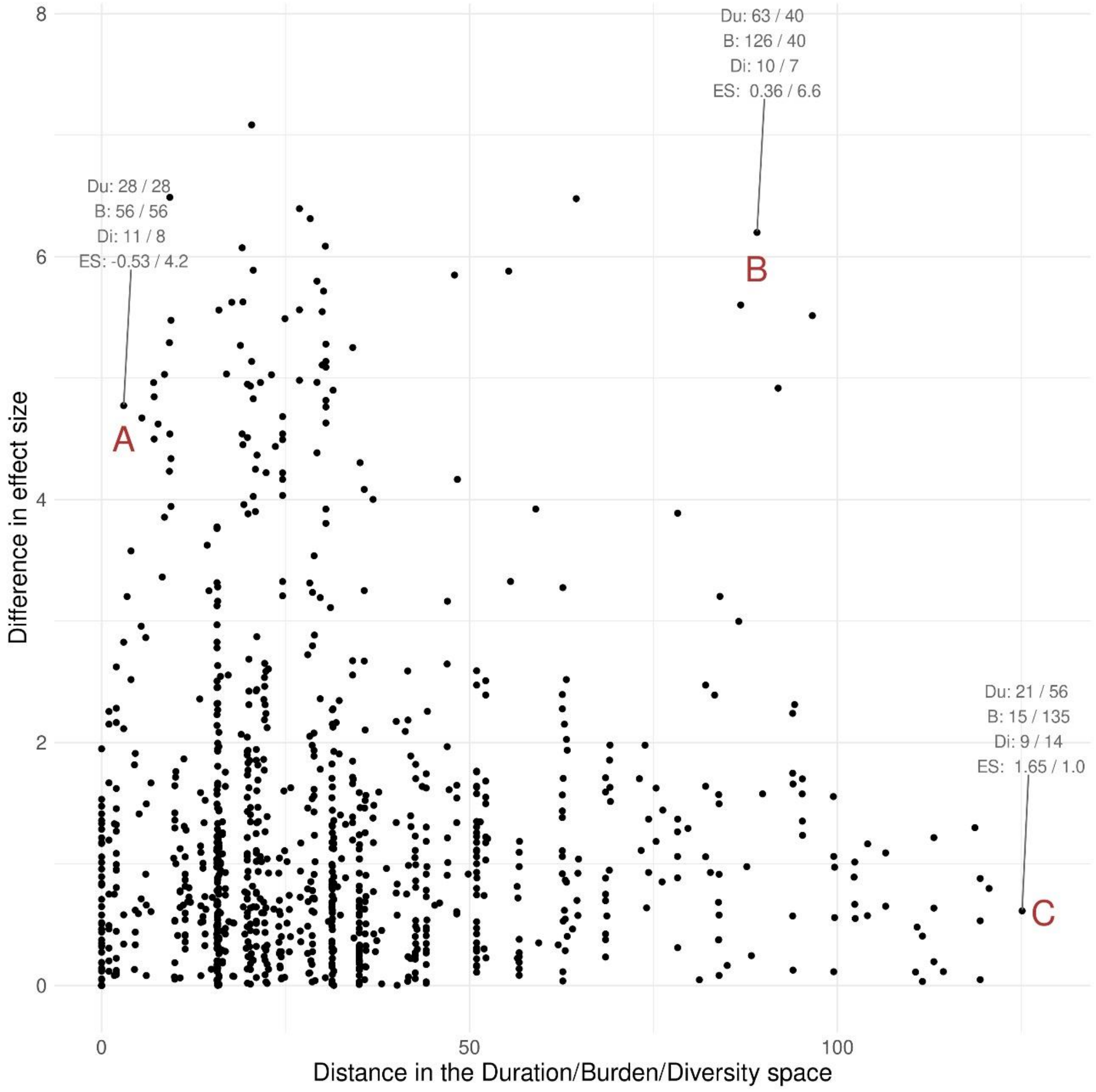


***Figure B: An annotated version of Figure 5 from the main text; correlation between the Euclidean distance of CVS characteristics and the difference in FST effect size.***

#### *3. Selecting an effect size to use in a sample size calculation*

In the main text, we recommend selecting an effect size of 2.0 (Cohen's d) as biologically meaningful for the sucrose preference test (SPT). We used the model shown here to justify this recommendation (Figure C). We observed a median sucrose preference of 79.9% in control groups in the mouse SPT, so we selected a control sucrose preference of 80% and modelled effect sizes for a range of reductions in post-CVS sucrose preference using different standard deviations (SD). As expected, effect size increased as the difference between the control and test means increased, and increasing the standard deviation reduced the effect size for a given difference between means. We judged a post-CVS sucrose preference of 60% to be biologically meaningful. We observed a median SD of 10.1% in control mouse SPTs and a median SD of 9.8%

in post-CVS mouse SPTs, so we selected an SD of 10% (the green line in Figure C) to arrive at the recommended effect size of 2.0.

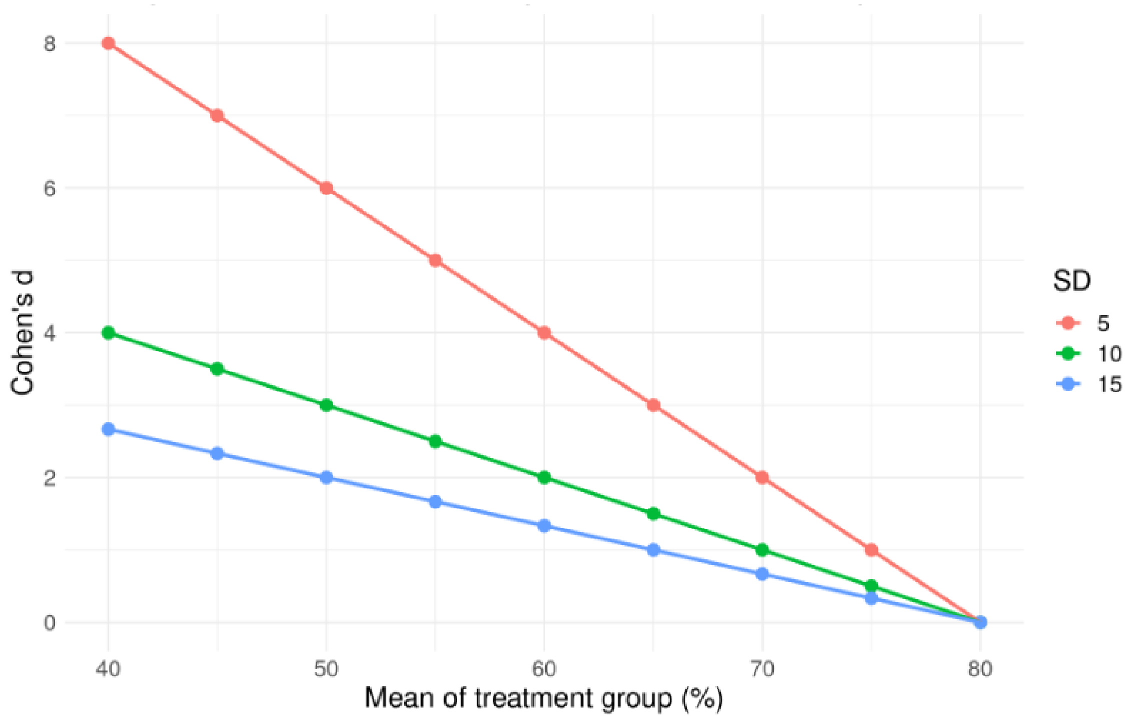


***Figure C: Change in effect size (Cohen's d) as a function of the difference between means, and variability of, control and test data. SD is standard deviation.***
