## Supplementary Tables 1-9 - SPT, TST, FST, OFT and EPM characteristics in mouse and rat studies for "Rodent chronic variable stress procedures: a disjunction between stress entity and impact on behaviour"

**Supplementary Table 1: Characteristics of the sucrose preference tests used in mouse studies.** If the article did not mention prior water or food deprivation, we assumed no deprivation took place and recording a duration of 0 h.

| **PubMed ID** | **Test duration (h)** | **% sucrose solution (w/v)** | **Duration of prior water deprivation (h)** | **Duration of prior food deprivation (h)** |
| --- | --- | --- | --- | --- |
| 30954529 | 1 | 1 | 24 | 24 |
| 32453814 | 1 | 1 | 24 | 24 |
| 30458370 | 2 | 1 | 0 | 0 |
| 34295228 | 2 | 2 | 18 | 18 |
| 32201112 | 3 | 2 | 24 | 0 |
| 37270930 | 3 | 2 | 24 | 0 |
| 37530136 | 3 | 2 | 24 | 24 |
| 37678448 | 3 | 2 | 24 | 24 |
| 35724494 | 4 | 1 | 0 | 0 |
| 30502341 | 6 | 1 | 24 | 24 |
| 37308778 | 6 | 1 | 24 | 24 |
| 31790664 | 12 | 1 | 0 | 0 |
| 36573299 | 12 | 1 | 24 | 0 |
| 36909192 | 12 | 1 | 12 | 0 |
| 36921537 | 12 | 1 | 24 | 24 |
| 37716030 | 12 | 1 | 0 | 24 |
| 31439031 | 24 | 1 | 0 | 0 |
| 32387133 | 24 | 1 | 0 | 0 |
| 32488052 | 24 | 1 | 0 | 0 |
| 35029057 | 24 | 1 | 0 | 0 |
| 36195262 | 24 | 1 | 0 | 0 |
| 36610138 | 24 | 1 | 0 | 0 |
| 37239235 | 24 | 2 | 12 | 0 |
| 37358676 | 24 | 2 | 24 | 0 |
| 35377087 | 24 | 1 | 0 | 12 |
| 32265492 | 24 | 2 | 0 | 0 |
| 33951503 | 48 | 1 | 0 | 0 |
| 37473999 | Not given | 1 | 0 | 0 |
| 35973368 | Not given | Not given | Not given | Not given |

**Supplementary Table 2: Characteristics of the tail suspension tests used in mouse studies.**

| **PubMed ID** | **Test duration (min)** | **Immobilisation measured for:** | **Mitigation for tail-climbing behaviour** |
| --- | --- | --- | --- |
| 35724494 | 4 | all 4 min | Unknown |
| 32265492 | 5 | all 5 min | Unknown |
| 35377087 | 6 | all 6 min | Unknown |
| 36909192 | 6 | all 6 min | Unknown |
| 31790664 | 6 | all 6 min | Cylinder used |
| 32488052 | 6 | all 6 min | Cylinder used |
| 30502341 | 6 | last 4 min | led to exclusion |
| 30954529 | 6 | last 4 min | led to exclusion |
| 32201112 | 6 | last 4 min | led to exclusion |
| 32453814 | 6 | last 4 min | led to exclusion |
| 31439031 | 6 | last 4 min | Unknown |
| 32387133 | 6 | last 4 min | Unknown |
| 34295228 | 6 | last 4 min | Unknown |
| 35029057 | 6 | last 4 min | Unknown |
| 36195262 | 6 | last 4 min | Unknown |
| 36921537 | 6 | last 4 min | Unknown |
| 37239235 | 6 | last 4 min | Unknown |
| 37308778 | 6 | last 4 min | Unknown |
| 37473999 | 6 | last 4 min | Unknown |
| 37530136 | 6 | last 4 min | Unknown |
| 37678448 | 6 | last 4 min | Unknown |
| 37716030 | 6 | last 4 min | Unknown |
| 37270930 | 6 | last 4 min | Cylinder used |
| 37358676 | 6 | last 4 min | Cylinder used |
| 33951503 | 6 | last 5 min | Unknown |
| 36573299 | 6 | last 5 min | Unknown |
| 30458370 | 6.5 | last 6 min | Unknown |
| 36610138 | 8 | last 6 min | Unknown |
| 35973368 | Not given | Not given | Not given |

**Supplementary Table 3: Characteristics of the forced swim tests used in mouse studies.**

| **PubMed ID** | **Test duration (min)** | **Immobilisation measured for:** |
| --- | --- | --- |
| 30458370 | 5 | all 5 min |
| 32265492 | 5 | all 5 min |
| 33951503 | 5 | last 4 min |
| 31790664 | 6 | all 6 min |
| 35377087 | 6 | all 6 min |
| 36909192 | 6 | all 6 min |
| 35184325 | 6 | all 6 min |
| 35664490 | 6 | all 6 min |
| 37091920 | 6 | all 6 min |
| 30502341 | 6 | last 4 min |
| 30954529 | 6 | last 4 min |
| 31439031 | 6 | last 4 min |
| 32201112 | 6 | last 4 min |
| 32387133 | 6 | last 4 min |
| 32453814 | 6 | last 4 min |
| 32488052 | 6 | last 4 min |
| 34295228 | 6 | last 4 min |
| 35029057 | 6 | last 4 min |
| 35724494 | 6 | last 4 min |
| 36195262 | 6 | last 4 min |
| 36610138 | 6 | last 4 min |
| 36921537 | 6 | last 4 min |
| 37239235 | 6 | last 4 min |
| 37270930 | 6 | last 4 min |
| 37308778 | 6 | last 4 min |
| 37358676 | 6 | last 4 min |
| 37473999 | 6 | last 4 min |
| 37530136 | 6 | last 4 min |
| 37678448 | 6 | last 4 min |
| 37716030 | 6 | last 4 min |
| 36704356 | 6 | last 4 min |
| 37091920 | 6 | last 4 min |
| 30856395 | 6 | last 4 min |
| 36573299 | 6 | last 5 min |
| 35973368 | Not given | Not given |

**Supplementary Table 4: Characteristics of the open field tests used in mouse studies.**

| **PubMed ID** | **Test duration (min)** |
| --- | --- |
| 30856395 | 5 |
| 33951503 | 5 |
| 33727046 | 5 |
| 36936941 | 5 |
| 37091920 | 8 |
| 32488052 | 10 |
| 35184325 | 10 |
| 35664490 | 10 |
| 36736870 | 10 |
| 36909192 | 10 |
| 31821847 | 10 |
| 33171149 | 10 |
| 35880047 | 10 |
| 36704356 | Not given |

**Supplementary Table 5: Characteristics of the elevated plus maze tests used in mouse studies.**

| **PubMed ID** | **Test duration (min)** |
| --- | --- |
| 30856395 | 5 |
| 33951503 | 5 |
| 35184325 | 5 |
| 37091920 | 5 |
| 33171149 | 5 |
| 33727046 | 5 |
| 35880047 | 5 |
| 36936941 | 5 |
| 35664490 | 6 |
| 36909192 | 6 |
| 32488052 | 10 |
| 36736870 | 10 |
| 31821847 | 10 |
| 36704356 | Not given |

**Supplementary Table 6: Characteristics of the sucrose preference tests used in rat studies.** If the article did not mention prior water or food deprivation, we assumed no deprivation took place and recording a duration of 0 h.

| **PubMed ID** | **Test duration (h)** | **% sucrose solution (w/v)** | **Duration of prior water deprivation (h)** | **Duration of prior food deprivation (h)** |
| --- | --- | --- | --- | --- |
| 33186383 | 1 | 1 | 0 | 22 |
| 33675839 | 1 | 1 | 4 | 4 |
| 31375779 | 1 | 1 | 4 | 0 |
| 30654121 | 1 | 1 | 4 | 0 |
| 30524234 | 1 | 1 | 4 | 0 |
| 34166749 | 1 | 1 | 12 | 0 |
| 32621862 | 1 | 1 | 12 | 0 |
| 35640844 | 1 | 1 | 14 | 14 |
| 37555927 | 1 | 1 | 23 | 23 |
| 33112507 | 1 | 1 | 23 | 0 |
| 37734259 | 1 | 1 | 24 | 24 |
| 37539173 | 1 | 1 | 24 | 24 |
| 36892979 | 1 | 1 | 24 | 24 |
| 36852042 | 1 | 1 | 24 | 24 |
| 34982171 | 1 | 1 | 24 | 0 |
| 31229467 | 1 | 1 | 24 | 24 |
| 35308874 | 1 | 1.5 | 18 | 18 |
| 34488064 | 1 | 2 | 0 | 0 |
| 34664178 | 1 | 2 | 16 | 0 |
| 34674723 | 3 | 1 | 24 | 24 |
| 31108115 | 4 | 1 | 1 | 1 |
| 30179639 | 4 | 1 | 23 | 0 |
| 33895271 | 4 | 2 | 12 | 0 |
| 37041107 | 12 | 1 | 12 | 12 |
| 32629607 | 12 | 1 | 0 | 0 |
| 35455459 | 48 | 1 | 0 | 0 |
| 35838106 | Not given | 1 | 0 | 0 |
| 34220431 | Not given | 1 | 24 | 0 |
| 37654512 | Not given | Not given | Not given | Not given |

**Supplementary Table 7: Characteristics of the forced swim tests used in rat studies.**

| **PubMed ID** | **Test duration (min)** | **Immobilisation measured for:** |
| --- | --- | --- |
| 36852042 | 5 | all 5 min |
| 35838106 | 5 | all 5 min |
| 35455459 | 5 | all 5 min |
| 35308874 | 5 | all 5 min |
| 34982171 | 5 | all 5 min |
| 34674723 | 5 | all 5 min |
| 34664178 | 5 | all 5 min |
| 33675839 | 5 | all 5 min |
| 33112507 | 5 | all 5 min |
| 31375779 | 5 | all 5 min |
| 31108115 | 5 | all 5 min |
| 30654121 | 5 | all 5 min |
| 30179639 | 5 | all 5 min |
| 30524234 | 5 | all 5 min |
| 37555927 | 6 | all 6 min |
| 34220431 | 6 | all 6 min |
| 32629607 | 6 | all 6 min |
| 37539173 | 6 | last 4 min |
| 37041107 | 6 | last 4 min |
| 36892979 | 6 | last 4 min |
| 34166749 | 6 | last 4 min |
| 33186383 | 6 | last 4 min |
| 32621862 | 6 | last 4 min |
| 31229467 | 6 | last 4 min |
| 37734259 | 6 | last 5 min |
| 33895271 | 6 | last 5 min |
| 35640844 | 10 | all 10 min |
| 34488064 | 15 | all 15 min |
| 37654512 | 15 | all 15 min |

**Supplementary Table 8: Characteristics of the open field tests used in rat studies.**

| **PubMed ID** | **Test duration (min)** |
| --- | --- |
| 37879231 | 5 |
| 35898162 | 5 |
| 35842198 | 5 |
| 35842198 | 5 |
| 35741632 | 5 |
| 34213722 | 5 |
| 34039946 | 5 |
| 33654382 | 5 |
| 33344727 | 5 |
| 33186383 | 5 |
| 32579979 | 5 |
| 31881185 | 5 |
| 31108115 | 5 |
| 30880220 | 5 |
| 30367959 | 5 |
| 30367959 | 5 |
| 29990678 | 5 |
| 29990678 | 5 |
| 37521489 | 10 |
| 33607579 | 10 |
| 33321142 | 10 |
| 33230530 | 10 |
| 31758000 | 10 |
| 34481787 | Not given |
| 33007392 | Not given |
| 32734320 | Not given |

**Supplementary Table 9: Characteristics of the elevated plus maze tests used in rat studies.**

| **PubMed ID** | **Test duration (min)** |
| --- | --- |
| 37879231 | 5 |
| 37521489 | 5 |
| 35898162 | 5 |
| 35842198 | 5 |
| 35842198 | 5 |
| 35741632 | 5 |
| 34213722 | 5 |
| 33654382 | 5 |
| 33607579 | 5 |
| 33230530 | 5 |
| 33186383 | 5 |
| 32579979 | 5 |
| 31881185 | 5 |
| 31758000 | 5 |
| 31108115 | 5 |
| 30880220 | 5 |
| 30367959 | 5 |
| 30367959 | 5 |
| 29990678 | 5 |
| 29990678 | 5 |
| 34481787 | 10 |
| 33321142 | 10 |
| 32734320 | 15 |
| 34039946 | Not given |
| 33344727 | Not given |
| 33007392 | Not given |
